# Deodorize with Bugs: Inoculation with black soldier fly larvae alters the microbiome and volatile organic compound profile of decomposing food waste

**DOI:** 10.1101/2022.10.10.511663

**Authors:** Rena Michishita, Masami Shimoda, Seiichi Furukawa, Takuya Uehara

## Abstract

The black soldier fly (BSF; *Hermetia illucens*) is used in sustainable processing of many types of organic waste. However, organic waste being decomposed by BSF produces strong odors, hindering more widespread application. The odor components and how they are produced have yet to be characterized. We found that digestion of food waste by BSF significantly alters the microbial flora, based on metagenomic analyses, and the odor components generated, as shown by thermal desorption gas chromatography mass spectrometry analysis. Inoculation with BSF significantly decreased production of volatile organic sulfur compounds (dimethyl disulfide and dimethyl trisulfide), which are known to be released during methionine and cysteine metabolism by *Lactobacillus* and *Enterococcus* bacteria. BSF inoculation significantly changed the abundance of *Lactobacillus* and *Enterococcus* and decreased microbial diversity overall. These findings may help in optimizing use of BSF for deodorization of composting food waste.

## Introduction

As the global population continues to grow, the amount of organic waste generated also continues to increase and must be managed. For example, global average food waste is estimated at 121 kg/capita/year, according to the report on organic food waste by the United Nations Environment Programme (2021), and SDG 12.3 aims to reduce this by half by 2030. Organic waste generated in cities is processed in three major ways: incineration, landfill, and composting (Nanda & Berruti 2021). However, these waste disposal methods can cause secondary pollution, such as generation of greenhouse gasses, odors, and contamination of groundwater. Use of insects (crickets, litter beetle, black soldier fly, etc.) has recently shown promise for improved decomposition of food waste (Ojha et al. 2020). In particular, the black soldier fly (BSF; *Hermetia illucens*, Fig. 1a) can feed on many types of waste, including food residues and livestock manure, and the adult flies can be used as an alternative source of protein in livestock feed (Barragan-Fonseca et al. 2017; Kawasaki et al. 2019). Use of BSF also contributes to reduction of greenhouse gas emissions, such as NH_3_ and CH_4_, from livestock manure (Chen et al. 2019). Many aspects of BSF biology are being investigated, such as chromosome-scale genome sequencing (Generalovic et al. 2020), CRISPR-Cas9 genome editing (Zhan et al. 2020), low-cost rearing (Nakamura et al. 2016), bioconversion rate (Surendra et al. 2020), and potential application to circular bioeconomy (Liu et al. 2022); the results of these studies may reveal ways to improve the suitability of BSF for industrial-scale applications.

**Figure 1.**
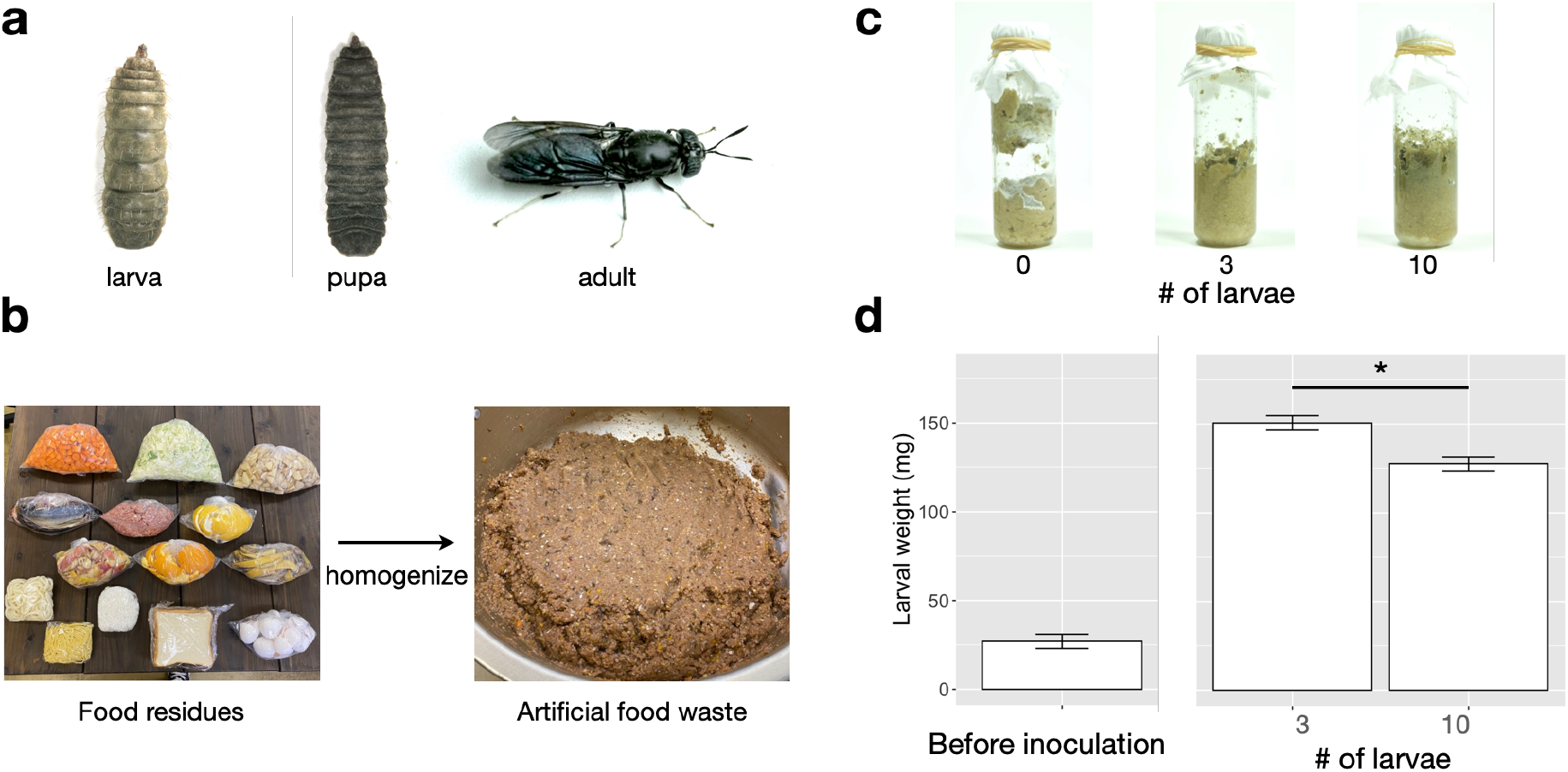
Black soldier fly (BSF) and methods for inoculating the larvae to the artificial food waste. **a,** Developmental stages of BSF. **b,** Schematic diagram of the method for preparing artificial food waste. **c,** Artificial food waste one week after BSF inoculation. **d,** Larval weights before and after BSF inoculation. Weights of larvae in the 3- and 10-larva vials were compared by using Student’s *t*-test. * significant difference (*P* < 0.05).

In most large-scale BSF breeding facilities, the waste containers and other facilities are open, and emit strong odors. On the other hand, when the flies are reared in closed, small-scale facilities, the odors decrease within a few days even if the flies are fed partially decomposed food scraps. In other words, BSF can decompose food waste without generating strong odors. Here, changes in the composition of volatile organic compounds (VOCs), especially those that generate odors, generated during processing of food scraps were identified by thermal desorption gas chromatography mass spectrometry (TDU-GC/MS) analysis. In addition, since the strong odors may be generated by microbial decomposition of food waste, we investigated how addition of BSF to decomposing food waste altered the microbial community by using metagenomic analysis.

## Results

### Digestion of organic waste by BSF larvae

The vials with and without BSF larvae differed in appearance. In no-larvae vials, only the upper surface of the food waste exposed to air was darker, and the food waste contained pockets of gas generated during decomposition (Fig. 1C). In vials with larvae, the food waste was uniformly black from exposure to the air, as the larvae mixed the scraps by moving through the vial while feeding. The vials with larvae did not contain pockets of gas.

The average weight of individual larva at inoculation was 26.96 ± 3.79 mg. After 7 days, the average weight of individual larva in 3- and 10-larva vials was 154.86 ± 5.46 mg and 129.85 ± 4.12 mg, respectively, increasing about 4–6 times in 7 days. The weight of larva reared in 3-larva vials was significantly higher than that of those reared in 10-larva vials.

### Alteration of VOCs by inoculation of BSF larvae

The number of odor components detected in the 0-, 3-, and 10-larva vials was 232.6 ± 27.1, 178.2 ± 6.2, and 182.0 ± 8.4, respectively (Fig. 2b,c). The three treatments did not differ significantly in the number of components, but the no-larvae treatment had greater variation in the number of components. In all vials, the most prominent peak was limonene, likely from fruit peels. In the principal component analysis of VOCs, each treatment clustered tightly in the principal component space (Fig. 2d).

**Figure 2.**
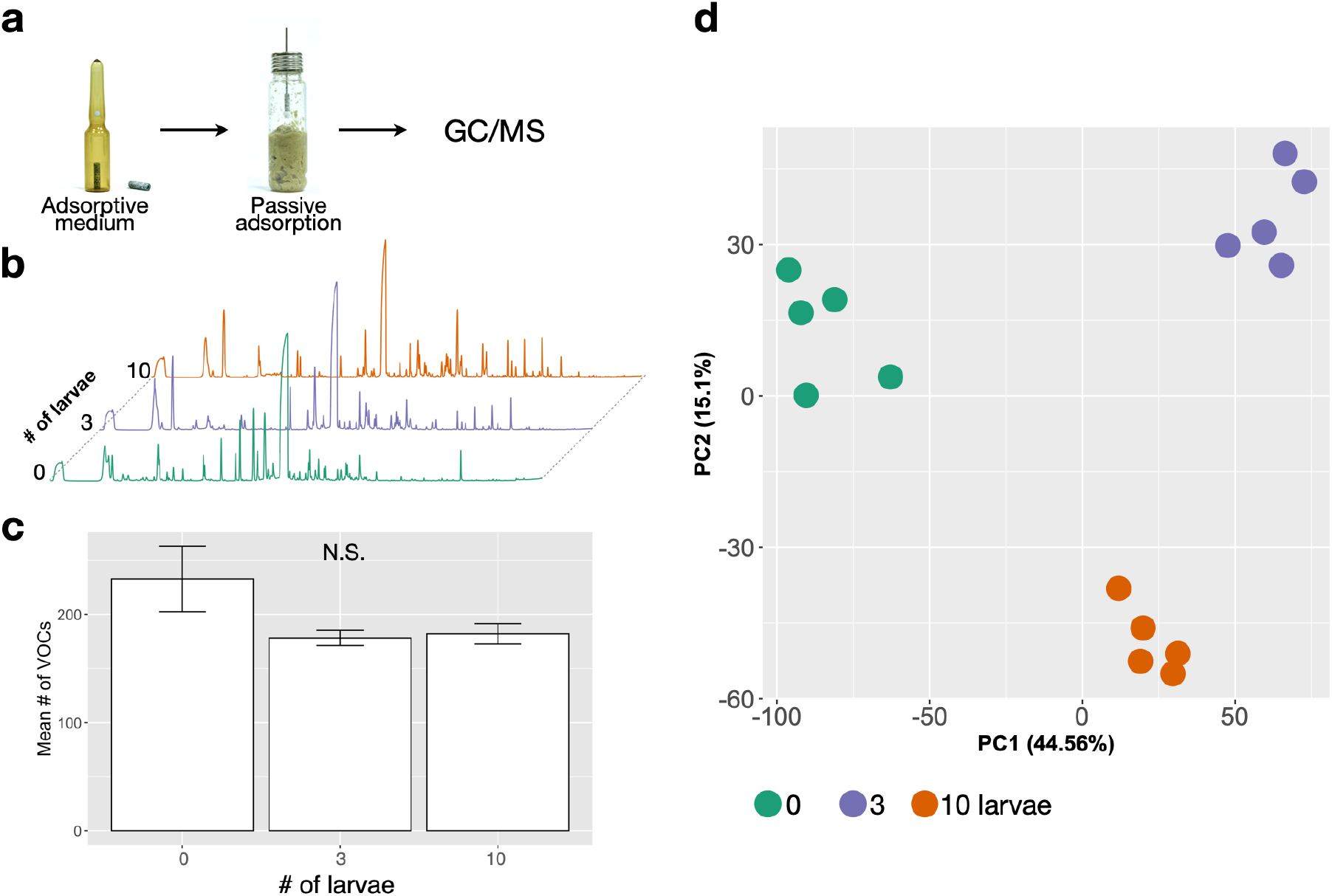
Overview of thermal desorption gas chromatography mass spectrometry (TDU-GC/MS) analysis of food waste. **a,** Adsorptive medium and its use for adsorption of odor components. **b,** Typical total ion chromatograms. **c,** Mean number of volatile organic compounds. Mean number of components did not differ significantly (N.S.) between treatments (ANOVA). **d,** Principal component (PC) analysis of odor components by treatment.

To investigate which odor components were altered by introduction of the BSF larvae, we performed a volcano plot analysis of components that differed between the 0- and 10-larva vials (Fig. 3a). Levels of 64 components were lower in the 10-larva vials than in the no-larvae vials (Supplementary Data 1). Some of these components are known to have foul odors, such as dimethyl disulfide (DMDS), dimethyl trisulfide (DMTS) and trimethylamine. In particular, comparison of DMDS and DMTS levels between the 0-, 3-, and 10-larva vials showed that the abundance of both components decreased with the addition of the BSF larvae (Fig. 3b).

**Figure 3.**
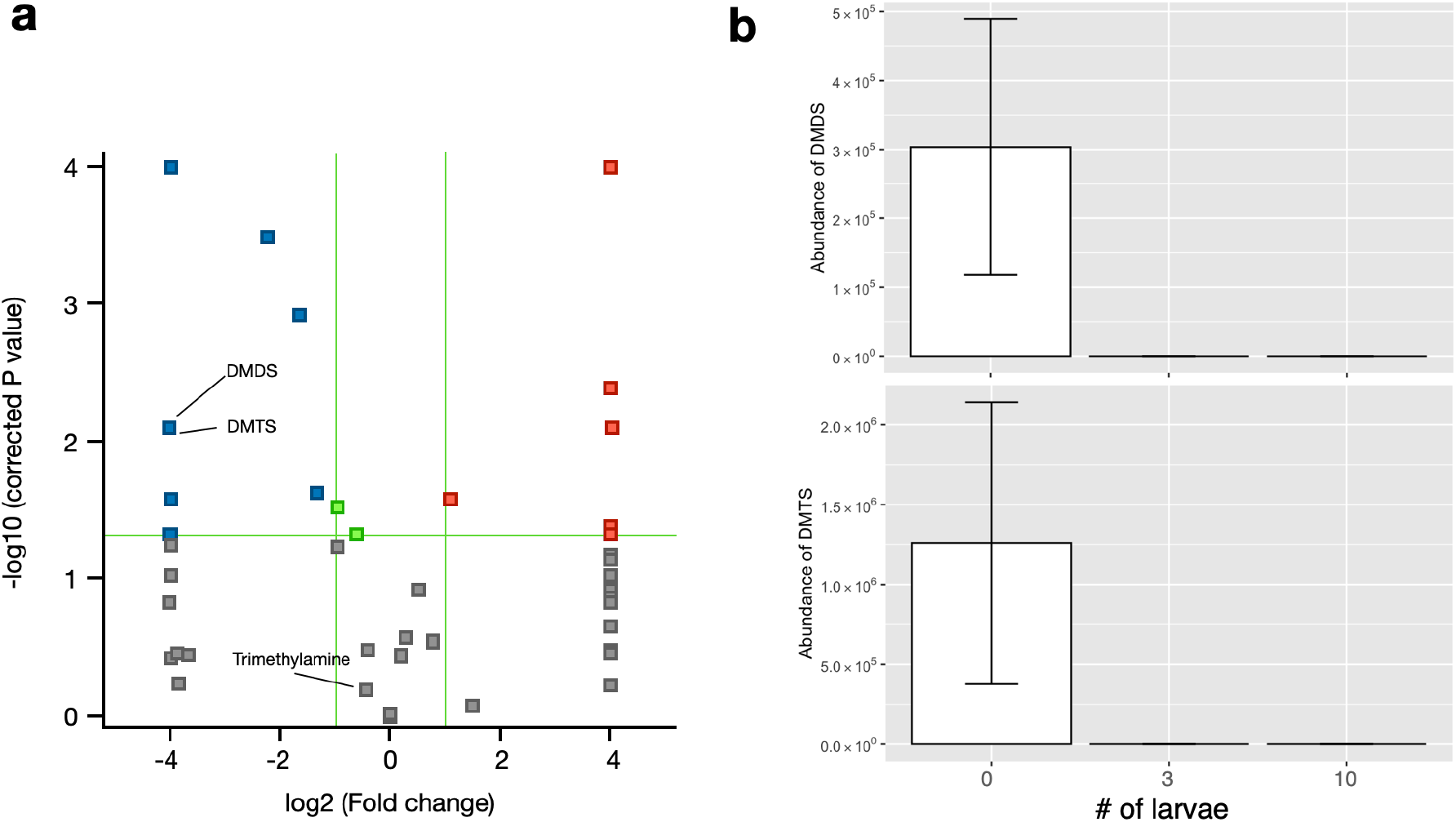
Odor components differentially detected between treatments. **a,** Volcano plot analysis of odor components detected in 0- and 10-larva vials. Blue and red squares show components detected with higher abundance in 0- and 10-larva vials, respectively. Green lines show thresholds of fold change and p-value. **b,** Abundance of DMDS (upper) and DMTS (lower) in each treatment.

To clarify the deodorizing effect of BSF larvae themselves, we added standard odor components to the artificial diet and compared odor profiles after 7 days. The amount of odor components was not significantly altered by BSF (Supplementary Fig. 1).

### Changes in the microbiome in food waste following inoculation of BSF larvae

We carried out metagenomic analysis of the microbes present in each treatment to assess how adding BSF to food waste changed the microbiome (Fig. 4a). The 16S rRNA gene PCR and MinION sequencing and analysis showed that most reads were assigned to 173 species and 34 genera (Supplementary Data 2, 3). To investigate the variation and qualitative differences in each treatment, we performed a non-metric multidimensional scaling (NMDS) analysis of species and number of reads. Data points for each of the three treatments clustered tightly in the NMDS space (Fig. 4b). Comparing alpha diversity in each treatment, microbial diversity decreased as the number of BSF larvae increased (Fig. 4c).

**Figure 4.**
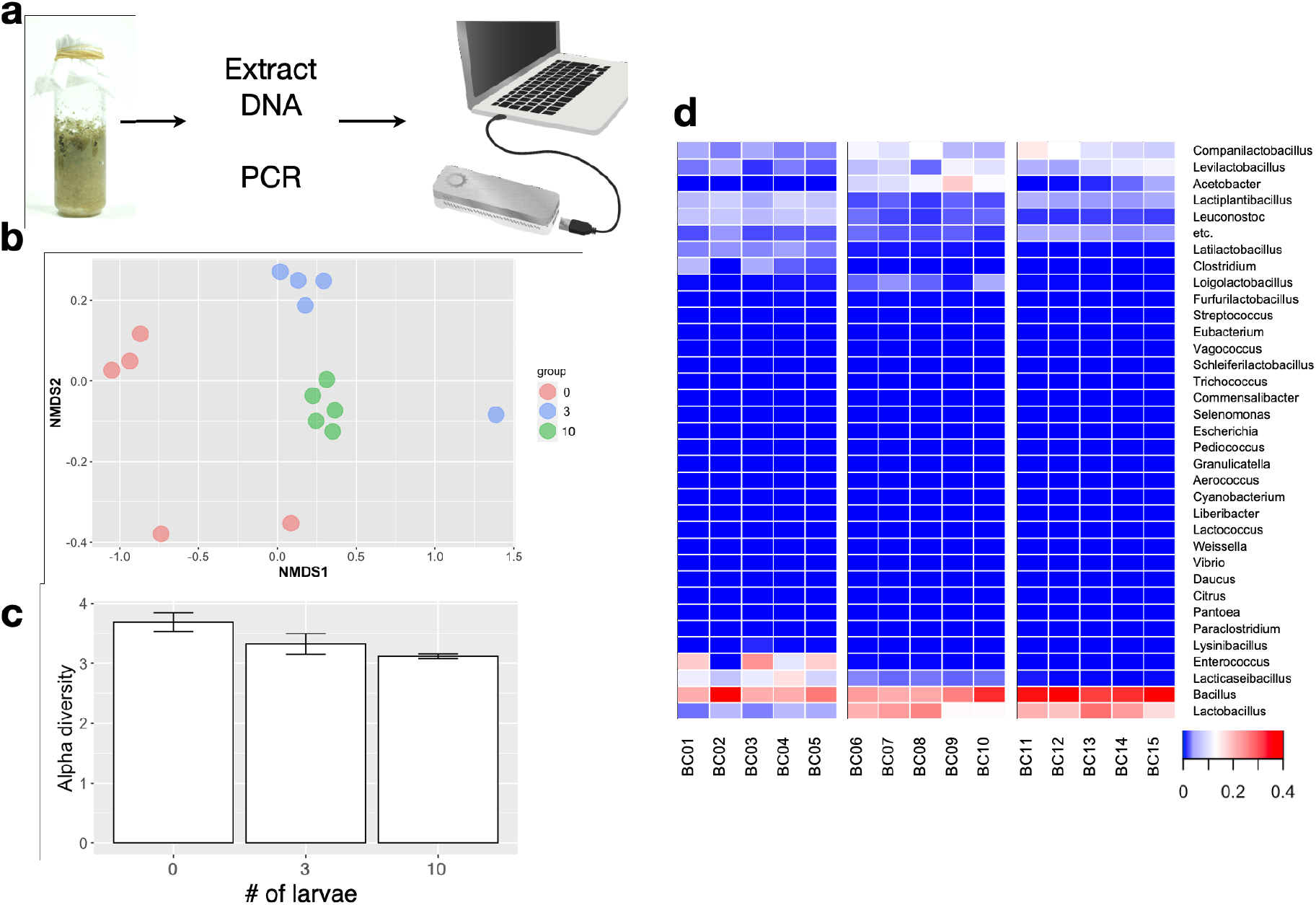
Overview of metagenomic analysis of microbes present in food waste. **a,** Schematic diagram of the methods for metagenomic analysis. **b,** Non-metric multidimensional scaling (NMDS) analysis of species detected by 16s amplicon sequencing. **c,** Comparison of alpha diversity (Shannon Index) of species detected in each treatment. **d**, Heatmap analysis of genera detected in each treatment (no-larvae: BC01–BC05, 3-larva: BC06–BC10, 10-larva: BC11–BC15).

Abundances of genera in each treatment were compared by heatmap analysis (Fig. 4d). *Bacillus* was the most abundant genus in every treatment. *Companilactobacillus, Levilactobacillus, Lactobacillus*, and *Acetobacter* were found only in vials with larvae. Additionally, abundances of *Bacillus, Companilactobacillus*, and *Levilactobacillus* increased as the number of larvae increased. On the other hand, *Enterococcus* and *Lacticaseibacillus* were observed only in the no-larvae vials. Most of the microbes identified are anaerobic, and there was no evidence that introduction of BSF favored the growth of either aerobic or anaerobic microorganisms.

## Discussion

Our TDU-GC/MS analysis showed that inoculating food waste with BSF larvae caused qualitative and quantitative changes in VOC composition, which may explain the differences in the odors produced when BSF larvae are used for decomposition of food waste. Metagenomic analysis revealed that introduction of BSF also significantly changed the composition of the microbial community in food waste. These results suggest that introduction of BSF larvae into food waste causes physical and biochemical changes in the degradation process, altering the microbial community and the odor composition.

### Physicochemical and biological effects on microorganisms

Most previous studies focused on the gut microbial flora of BSF larvae (Klammsteiner et al. 2020; 2021; Zhan et al. 2020; Zhineng et al. 2021), and research focusing on the surrounding microbiological environment is extremely limited. Studies on the BSF gut microbial flora identified *Morganella* and *Dysgonomonas* as genera that are typically present (Klammsteiner et al. 2020; Tanga et al. 2021; Zhan et al. 2020); however, we did not find these bacteria in our analyses. We found that introduction of BSF larvae to organic waste resulted in decreased alpha diversity, or species richness, of the microbial flora; microbial diversity decreased as the number of larvae increased. Wu et al. (2021) also reported lower microbial species diversity in larval frass, which is consistent with our results. The observed changes in microbial flora seem to be due to physical and biochemical changes caused by the larvae, and to depend on the number of larvae introduced, as described below.

The primary effect of introducing BSF larvae into food waste is physical agitation of the waste as the larvae move through it by peristaltic action. We found that waste with and without larvae showed distinct physical differences (Fig. 1c). In no-larvae vials, the gas produced by decomposition created pockets in the food waste. Vials with larvae lacked these gas pockets because the agitation of the food waste by the larvae allowed the gas to escape to the vial headspace. Food waste in vials with larvae turned black, likely due to oxidation of polyphenols contained in the fruit peels. In addition, the larvae were able to digest and metabolize available nutrients in the waste and to grow to maturity. Although the growth rate of larvae varied with the age, larval weight increased by a factor of approximately 4 or more in 7 days in our experiments. Bioconversion rates and physicochemical changes in organic wastes following introduction of BSF larvae have been reported in several studies (Klammsteiner et al. 2021, Li et al. 2022). In general, digestion by BSF larvae tends to make the pH of waste more basic (pH 8.0–9.0, Ma et al. 2018) and to decrease the nutrient content (Klammsteiner et al. 2021). These changes are likely to have a significant impact on the microbial community in the food waste.

BSF larvae can grow in environments with high levels of pathogenic microorganisms, such as livestock manure, and are thought to have a highly developed immune system. The BSF genome contains 50 genes that encode antimicrobial peptides, one of the largest numbers so far identified in insects (Zhan et al. 2020). Several peptides, such as defencin, diptericin, and stomoxyn, have been cloned from BSF and confirmed to show antimicrobial activity in in vitro experiments (Elhag et al. 2017; Xu et al. 2020). A transgenic silkworm expressing these three genes showed an increased resistance to both gram-positive and gram-negative entomopathogenic bacteria (Xu et al. 2020). These findings suggest that not only physical but also biochemical effects, such as production of immunity-related compounds, are involved in the alteration of the microbial flora.

### Effects on odor composition

Inoculation with BSF larvae changed the composition of odor components. We hypothesize that these changes reflect changes in the microbiota and the odor compounds they produce. Although information on the relationship between VOC emission and microbial flora is extremely limited, several bacteria are reported to affect emission of VOCs. For example, volatile organic sulfur compounds (VOSCs) such as DMDS and DMTS are positively correlated with the presence of microorganisms including *Lactobacillus* and *Leuconostoc* (Zhang et al. 2020). Another study showed that *Bacillus* and *Actinomycetes* might produce VOSCs during decomposition (Mayrhofer et al. 2006). VOSCs are known as key components from meat and vegetable wastes (Zhang et al. 2020). During catabolism of sulfur-containing amino acids such as methionine and cysteine, microorganisms including lactic acid bacteria and *Bacillus* produce methanethiol, which can easily be converted to DMDS and DMTS (Hanniffy et al. 2009, Seefeldt & Weimer 2000). Conversion of these amino acids is catalyzed by microbial enzymes, and the enzymatic activity differs among strains (Hanniffy et al. 2009). We found that the abundance of both VOSCs and *Lactobacillus, Leuconostoc, Lacticaseibacillus*, and *Enterococcus* differed significantly between BSF-inoculated and control vials, which is consistent with these reports.

Although BSF larvae appear to reduce VOSC emission via alteration of the microbial flora, BSF larvae themselves seem to have less ability to reduce these components directly (Supplementary Fig. 1). For example, there was no difference in levels of limonene, which is contained in fruit peels, between BSF-inoculated and control vials. In other words, the lower the initial odor of the organic waste, the higher the deodorizing effect.

### Potential use of BSF to control odors in organic waste recycling

In general, organic waste processing with BSF is known to produce strong odors. Our results, however, showed that the introduction of BSF can reduce certain odor components. BSF processing is also likely to be effective for reducing the odor of livestock manure (Beskin et al. 2018), and we believe that our results can be generalized not only to food waste but also to other organic waste. Below we present our recommendations for eliminating odors during processing of organic waste with BSF. Our results show that the deodorizing effect of the BSF itself is small. Therefore, it is unlikely that BSF can reduce the odor of organic waste that already has a strong odor by feeding on it. On the other hand, if the odor of the organic waste is caused by microbial decomposition, introduction of BSF at the early stage can be highly effective in eliminating odors. The insect’s ability to dramatically alter the surrounding microbiological environment would strongly inhibit the generation of odorous components such as DMDS and DMTS. To minimize the foul odors and maximize production of protein in the form of adult flies, it is important to have a high waste to larvae ratio. However, to optimize deodorization, a lower waste to larvae ratio is preferable to maximize the biochemical and physical effects.

## Methods

### Insects

Successive generations of lab-reared flies were used in this study. Flies were originally collected in Tsukuba, Japan in 2013, fed an artificial diet for the vinegar fly, and maintained in plastic containers (12 cm diameter, 10 cm height, Mineron-Kasei Co., Ltd., Osaka, Japan) under a 16L:8D photoperiod, at 25°C (Nakamura et al. 2016).

### Artificial food waste

Experimental designs for BSF research vary widely from study to study, making it difficult to compare results between studies (Bosch et al. 2020). We here standardized an artificial food waste in the lab (Fig. 1b), based on the food found in a typical Japanese home, using food from the following five categories: vegetables, fruits, carbohydrates, meat, and fish (Supplementary Table 1; Food Waste Suitable for Treatment Using Black Soldier Fly Larvae, Hirayasu et al. 2017). To create decomposed waste prior to addition of larvae, freshly prepared waste was kept for 3 days in the same conditions used for rearing the BSF. Decomposing food waste (10 g) was dispensed into a glass vial (2.12 cm diameter, 7.55 cm height, Gerstel GmbH and Co., KG, Germany). Then 3 or 5 individuals of 10-day old larvae were washed with distilled water and transferred to the vial. Vials without larvae were used as controls.

### VOC sampling

Seven days after larval inoculation, volatile components in the vial headspace were collected by passive adsorption onto a Monotrap (Fig. 2a, RGC18 TD, GL-Science, Tokyo, Japan) for 3 h. Following adsorption, samples were transferred to a 1.5-ml vial and stored at −20°C until analysis.

### TDU-GC/MS analysis

Headspace volatiles collected using the Monotrap were analyzed by gas chromatography-mass spectrometry (GC-MS, GC: Agilent 7890A/MS5977B MSD, Agilent Technologies, CA, USA) with an HP-5MS UI capillary column (30 m, 0.25-mm ID, 0.25-μm film thickness; Agilent Technologies) equipped with a thermo-desorption system, cooled injection, and cold trap (Gerstel GmbH and Co.). The GC was maintained at 40°C for 3 min, increased to 150°C at a rate of 10°C/min, then to 280°C at 20°C/min, and held at that temperature for 5 min. Helium was the carrier gas at a constant flow of 1.1 ml/min. The compounds were tentatively identified using data contained in the NIST Mass Spectral Library, 2017 release.

### Data analysis

GC/MS data were deconvoluted using Unknowns Analysis software (ver. B.09.00, Agilent Technologies Inc.) and aligned using Mass Profinder Professional (ver. 14.9, Agilent Technologies Inc.). Peaks with amplitudes of less than 1% of the maximum peak height were ignored. Only entities present in at least 60% of replicates from one condition were included in subsequent analyses. To determine changes in components in each condition, the data were subjected to principal component analysis by using the statistical software R (version 4.1.0, R Core Team 2021). To determine which components varied between conditions, we performed volcano plot analysis on the 0- and 10-individual vials (*P*-value 0.05, fold change 2) in the Mass Profinder Professional software. Compound identification was performed by comparing mass spectra to the NIST 2017 Library.

### Metagenome analysis

DNA was extracted from the rotten food waste with NucleoSpin DNA Stool kits (Macherey-Nagel GmbH & Co. KG, Düren, Germany) and quality checked by using a NanoDrop (Thermo Fisher Scientific Inc., MA, USA) and Qubit 4 Fluorometer (Thermo Fisher Scientific Inc.). Sequencing libraries were prepared with a 16S Barcoding Kit (SQK-16S024, Oxford Nanopore Technologies, UK) and sequenced by using MinION (Oxford Nanopore Technologies). The data were demultiplexed by using the adapter trimming tool Porechop (Wick et al. 2017), and Emu (Curry et al. 2022) was used to estimate sequence relative abundance, with default settings. NMDS analysis was performed by using the *vegan* package in R (Oksanen et al. 2020).

## Data availability

All raw reads have been uploaded to DDBJ under BioProjectID PRJDB14354 and SRA under DRA014852.

## Contributions

R.M., M.S., and T.U. conceived the project and designed and interpreted all of the experiments. R.M. performed the experiments shown in Figs. 1–4. M.S. helped to design and perform insect conditioning and artificial food waste preparation. T.U. analyzed data for all figures and helped to perform TDU-GC/MS and 16s amplicon analyses. R.M., M.S., and T.U. wrote the paper with help from S.F.

## Acknowledgements

We thank Drs. Tetsuya Kobayashi and Liu Chia-Ming for comments on the manuscript, and members of the Insect Design Technology Group for maintenance of the BSF colony. This work was supported by the Cabinet Office, Government of Japan, Cross-ministerial Moonshot Agriculture, Forestry and Fisheries Research and Development Program, “Technologies for Smart Bio-industry and Agriculture” (funding agency: Bio-oriented Technology Research Advancement Institution) [JPJ009237]. This research was in partial fulfillment of an MSc degree (RM) from the University of Tsukuba. The MinION image in Fig. 4a was provided by DBCLS Togo Picture Gallery (©2016 DBCLS TogoTV; https://togotv.dbcls.jp/pics.html).

## Supplementary Information

**Supplementary Table 1.** Contents of artificial food waste.

| Category | Content | Weight (g) |
| --- | --- | --- |
| Vegetables | Cabbage | 170 |
|  | Carrot | 170 |
|  | Potato | 160 |
| Fruits | Apple peel | 50 |
|  | Banana peel | 50 |
|  | Orange peel | 40 |
|  | Grapefruit peel | 40 |
| Animal proteins | Minced pork | 80 |
|  | Small horse mackerel | 100 |
|  | Egg shell | 20 |
| Carbohydrates | Rice | 30 |
|  | Bread | 30 |
|  | Udon noodles | 30 |
|  | Chinese noodles | 30 |

**Supplementary Figure 1.**
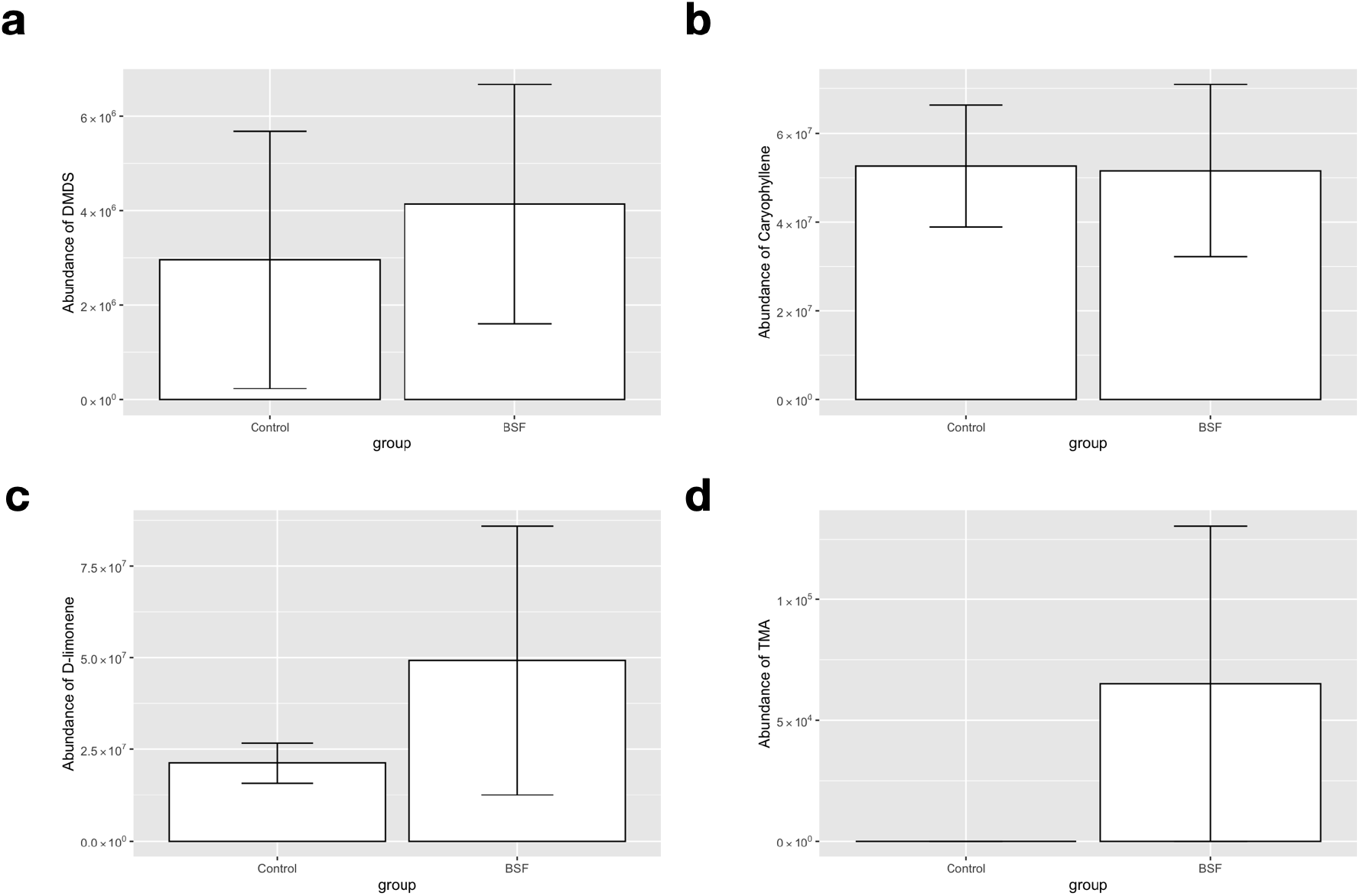
Abundance of odor components 7 days after black soldier fly (BSF) introduction. Abundances of (**a**) dimethyl disulfide (DMDS), (**b**) caryophyllene, (**c**) D-limonene and (**d**) trimethylamine. To clarify the deodorizing effect of BSF larvae themselves, we added standard odor components to the artificial diet and compared the residual odor components after 7 days. Components were detected and analyzed by GC/MS as mentioned above. The following chemicals were used as odor standards: DMDS, D-limonene, beta-caryophyllene (TCI Chemical Industry Co., LTD., Tokyo, Japan) and trimethylamine (FUJIFILM Wako Pure Chemical Industries, Ltd., Osaka, Japan).

